# Evolution of interface binding strengths in simplified model of protein quaternary structure

**DOI:** 10.1101/557272

**Authors:** Alexander S. Leonard, Sebastian E. Ahnert

**Affiliations:** Theory of Condensed Matter, Cavendish Laboratory, University of Cambridge, JJ Thomson Avenue, Cambridge CB3 0HE, United Kingdom; Sainsbury Laboratory, University of Cambridge, Bateman Street, Cambridge CB2 1LR, United Kingdom

## Abstract

The self-assembly of proteins into protein quaternary structures is of fundamental importance to many biological processes, and protein misassembly is responsible for a wide range of proteopathic diseases. In recent years, abstract lattice models of protein self-assembly have been used to simulate the evolution and assembly of protein quaternary structure, and to provide a tractable way to study the genotype-phenotype map of such systems. Here we generalize these models by representing the interfaces as mutable binary strings. This simple change enables us to model the evolution of interface strengths, interface symmetry, and deterministic assembly pathways. Using the generalized model we are able to reproduce two important results established for real protein complexes: The first is that protein assembly pathways are under evolutionary selection to minimize misassembly. The second is that the assembly pathway of a complex mirrors its evolutionary history, and that both can be derived from the relative strengths of interfaces. These results demonstrate that the generalized lattice model offers a powerful new framework for the study of protein self-assembly processes and their evolution.

## Author summary

Protein complexes assemble by joining individual proteins together through interacting binding sites. Because of the long time scales of biological evolution, it can be difficult to reconstruct how these interactions change over time. We use simplified representations of proteins to simulate the evolution of these complexes on a computer. In some cases the order in which the complex assembles is crucial. We show that biological evolution increases the strength of interactions that must occur earlier, and decreases the strength of later interactions. Similar knowledge of interactions being preferred to be stronger or weaker can also help to predict the evolutionary ancestry of a complex. While these simulations are not realistic enough to make exact predictions, this general link between ordered pathways in assembly and evolution matches well-established observations that have been made in real protein complexes. This means that our model provides a powerful framework for the study of protein complex assembly and evolution.

## Introduction

Many proteins self-assemble into protein quaternary structures, which fulfill a multitude of functions across a wide range of biological processes [1]. Abstract models trade off the complexity arising from conformations, buried surfaces, cooperative binding, etc., but still retain qualitative realism. A general class of polyomino tile self-assembly models have strong analytic potential while maintaining semblance to protein quatenary structure.

The polyomino self-assembly model [2] combines lattice tile self-assembly with a quantification of biological complexity, examining the relationship between genetic description length and phenotypic complexity. The same model was developed and expanded with evolutionary dynamics by Johnston *et al.* [3], and used to probe general properties of genotype-phenotype maps by Greenbury *et al.* [4].

Here we develop a generalization of interactions using binary strings in these polyomino assembly models, in particular introducing variable binding strengths and relaxing the rejection of misassembly.

Binding affinity is difficult to assess experimentally but central to making predictions on assembly [5]. A dominant cause in altering the affinity is mutations to polar or charged groups [6]. While our binary interface polyomino self-assembly model does not account for the variety of amino acids and their particular properties, it provides a reasonable coarse-grained approach. Similar models of protein interactions using binary subunit interfaces have linked protein-protein interaction properties to experimental observations on protein family evolution [7, 8].

Adding these features into polyomino models enables preliminary explorations into the evolution of binding strengths and the implications binding strengths can have on preferred evolutionary pathways.

Several recent studies have revealed the deep relationship between evolutionary pathways and assembly properties like stoichiometry [9], symmetry [10], interaction topology [1], and binding strengths [11]. We aim to reproduce several of these observations in the framework of our generalized polyomino model in order to highlight its potential as a tool for the study of protein assembly and its evolution.

### Self-assembly algorithm

Any self-assembling system requires two ingredients: assembly subunits with binding sites, and a method for determining the strength of an interaction between two such sites. The arrangement of the sites and their interactions can be described in the form of an assembly graph [12]. From these simple components, structures can be formed through the following stochastic assembly process:

- The process starts with a randomly chosen initial subunit.
- The structure grows by placing a randomly chosen subunit with random orientation in a random adjacent position to the existing structure.
- If the interaction interface between adjacent binding sites is sufficiently strong, the placed subunit binds irreversibly to the existing structure.
- The growth process repeats until no further bindings are possible. At this stage, assembly terminates and the final structure forms a single connected set of one or more subunits.

If the subunits are square tiles on a lattice, connected sets of tiles are called Polyominoes [13].

### Genotypes and Phenotypes

We can define a *genotype* that encodes a set of subunit interactions as a sequence, in which each sequence position represents the type of a particular binding site on a subunit. The assembly process maps a given genotype to a single polyomino (in the case of a deterministic genotype) or a statistical distribution of several different polyominoes (for a nondeterministic genotype). In either case these polyominoes can be thought of as abstract biological *phenotypes*.

The assembly process is independent of the order in which the subunits are represented in the genotype, and translations, rotations, or reflections of a given polyomino are not considered unique. The implementation of this invariance is outlined in S1 Appendix.

An example of the mapping from genotype to phenotype is shown in Fig 1, using the integer binding site conventions of existing polyomino models. Certain binding sites are noninteracting (labeled 0) while interactions of equal strength occur between fixed pairs of positive integers. The interacting pairs are 1 ↔ 2, 3 ↔ 4, etc.

**Fig 1.**
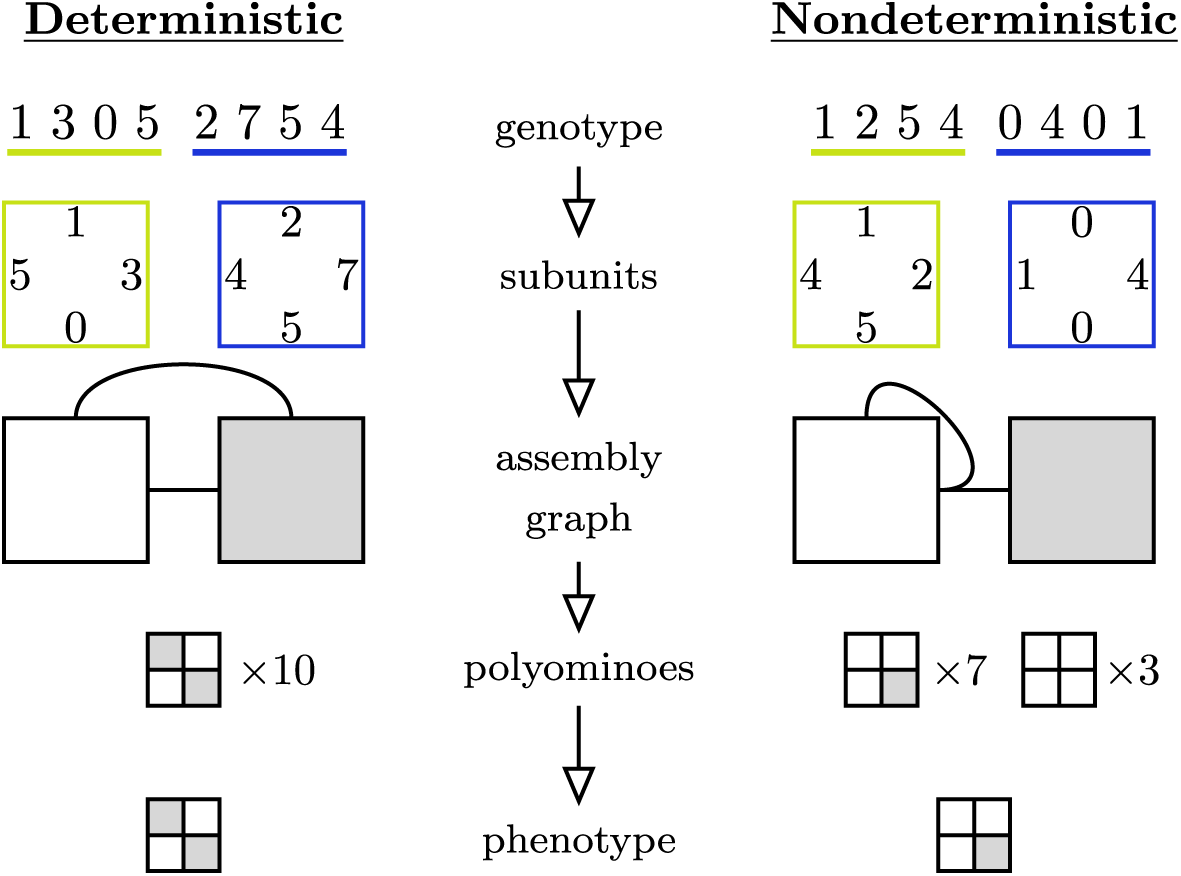
Assembly sequence from genotype to phenotype in the standard Polyomino self-assembly model. The full sequence of generating a phenotype from a genotype for deterministic (left) and nondeterministic (right) assemblies. The binding sites on the subunits are transcribed from the genotype in a clockwise fashion. The assembly graph encodes all possible interactions (0s noninteracting, 1s and 2s interact with each other, 3s and 4s interact with each other, etc.) among the subunits, indicated by solid lines. In the case of nondeterministic genotypes, different polyominoes may emerge as the outcomes of the stochastic assembly process. Here we perform 10 repeated assemblies, and define the phenotype of a genotype as the polyomino that appears most often. Other definitions of a phenotype from the distribution of polyominoes are also possible.

### Nondeterminism

Repeated assemblies of the same genotype do not necessarily produce the same polyomino, a property referred to as *nondeterminism*. There are many sources of nondeterminism, ranging from unbound aggregations of subunits to branching pathways in the course of the assembly process. A more general insight into nondeterminism in polyomino self-assembly is given by Tesoro, Ahnert, and Leonard [12].

Deterministic genotypes are significantly outnumbered by nondeterministic ones, and the addition of interactions typically increases the fraction of nondeterministic genotypes. In a biological context nondeterministic genotypes can be viewed as less desirable than deterministic ones, as the functions of many proteins strongly rely on the accuracy and reproducibility of their structures. We can therefore use nondeterminism in the polyomino self-assembly model to represent protein misassembly and thereby study the conditions under which proteins may evolve towards more stable and reliable assemblies.

### Generalized model framework

In this paper we generalize the standard Polyomino self-assembly model as outlined above by introducing interfaces that take the form of binary strings rather than integers. This definition of interfaces gives rise to further definitions of interface strength and symmetry. It also allows for non-transitive interactions between interfaces.

The assembly process outlined earlier is unchanged, with only the sites and thus how to determine interactions between them being redefined, as seen in Fig 2.

**Fig 2.**
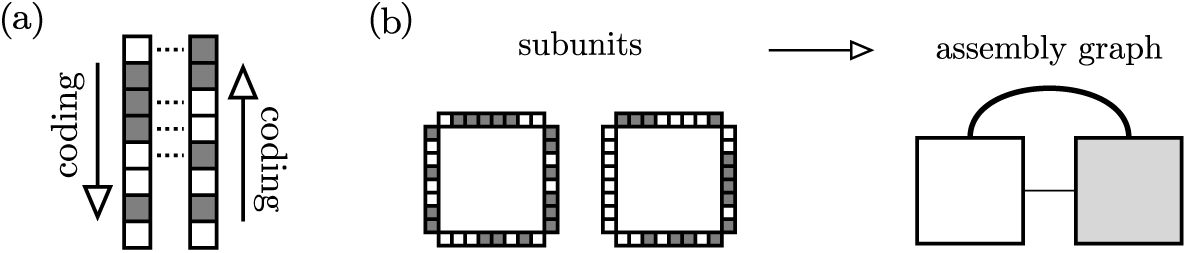
Generalized binding sites. (a) Explicit subsite interactions (dotted lines) between two binding sites, showing the “head to tail” alignment. The Hamming distance between the counter-aligned sites is 4, and so the interaction strength is *Ŝ* = .5. (b) Taking the critical strength *Ŝ*_*c*_ = .75, these two subunits encode two interactions in the assembly graph. The interactions have different strengths (indicated by line thickness), with the upper interaction stronger (*Ŝ* = .875) than the lower (*Ŝ* = .75).

The number of bits per binding site is given by *L*_*I*_, providing 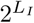 unique binding site configurations. Since the subunits are always encoded in a genotype following a common convention (e.g. clockwise around a tile), two adjoined sites have a “head to tail” alignment (see Fig. 2).

The interaction strength between two sites relates to the Hamming distance *d*_*H*_ between one site and the reversed alignment of the other, normalized by *L*_*I*_. As such, the interaction strength *Ŝ* ∈ [0, 1], and binding can occur if the strength is above some chosen critical strength *Ŝ* ≥ *Ŝ*_*c*_. The stochastic assembly process as outlined above is now extended to include a binding probability as a function of interaction strengths. Interacting subunits are no longer guaranteed to bind, but binding that does occur remains irreversible.

Binding probability can be linked to interaction strength via an abstract temperature *T* ∈ [0, ∞). More complex forms may have more physical justification, but a useful form of binding probability is

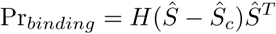

where *H* is the Heaviside function, taking *H*(0) = 1. The average number of attempts an interaction will take, effectively the binding time, is the reciprocal of the binding probability. With the choice *T* > 0, stronger bonds are expected to assemble more quickly than weaker bonds.

## Results

Using this model, even a small number of subunits can give rise to a large array of potential Polyomino structures. We focused our attention on a subset of six assembly graphs that contained both deterministic and nondeterministic phenotypes and transitions, and in which each of the four more complex assembly graphs are in principle accessible from two other members of the set via point mutations. The assembly graphs and phenotypes are shown in Fig 3.

**Fig 3.**
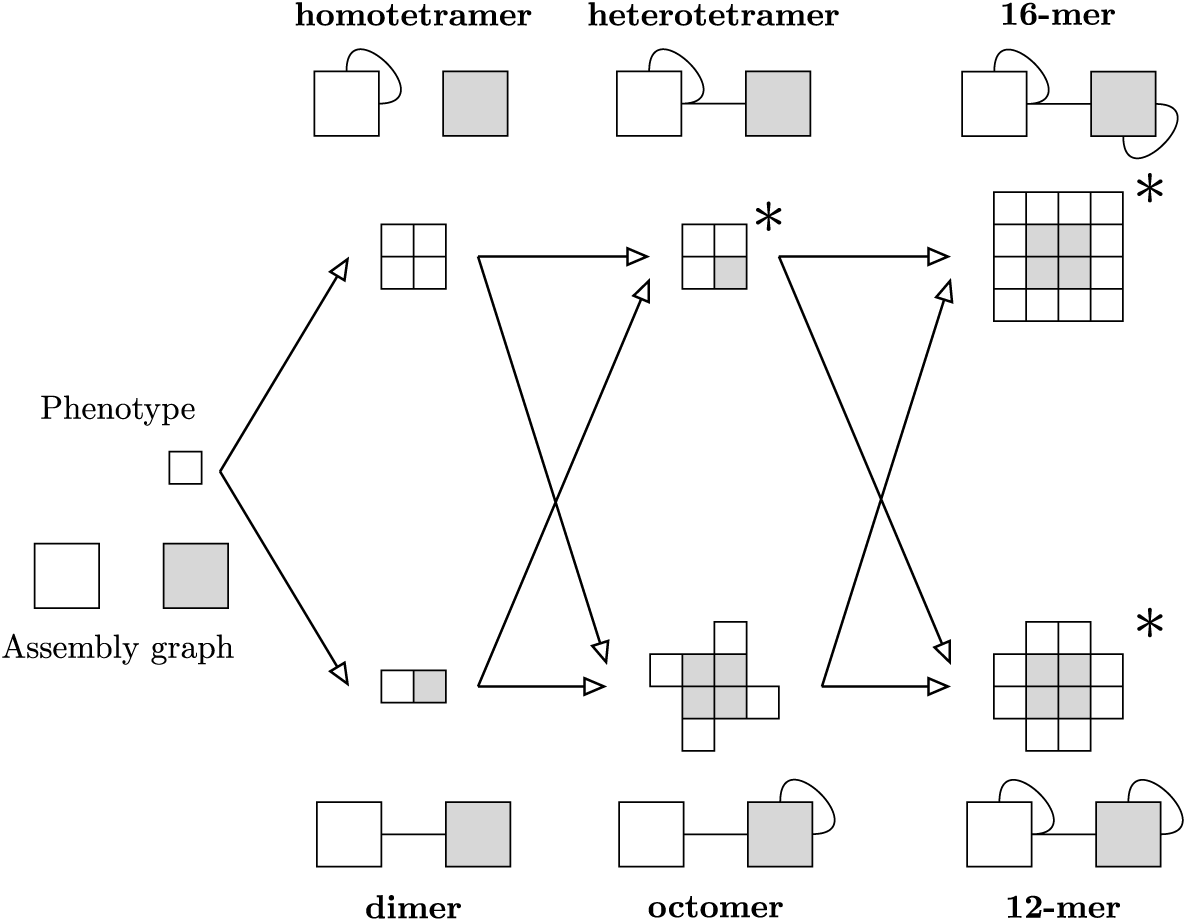
Example system of six assembly graphs. The interactionless initial condition and an example system of six assembly graphs with associated polyominoes. The assembly graphs (and polyominoes) are grouped into vertical columns that are ordered by the number of interactions (from left to right: one, two, and three interactions). Three assemblies are nondeterministic, and are marked with a ∗. In the nondeterministic cases we only show the most common Polyomino structure, which also corresponds to our formal definition of the phenotype.

Evolution was modeled with a fixed-size haploid population undergoing discrete generations of selection and mutation. Reproduction was asexual, and mutations occurred with a fixed probability to flip each bit in a genotype. Non-negative fitnesses were assigned to every individual according to their phenotype properties, with more fit members proportionally more likely to reproduce into the next generation. Nondeterminism was punished by an individual only receiving a fraction of its potential fitness equal to the frequency of correct assembly exponentiated by a parameter *γ* ∈ ℝ^≥1^.

## Binding strength dynamics

Accessing information on the evolution of real protein binding strengths over sufficiently long time scales is effectively impossible. There are potential proxies, like looking at homologous proteins across an evolutionary tree [14]. Experimental work has suggested a link between ordered assembly pathways and the constraints they place on evolution [11], but focused on subunits fusing together rather than individual strengths evolving.

Here we show how the generalized polyomino model can simulate evolutionary selection for assembly order, such as observed in [11] for real protein complexes. The possibility of nondeterminism in our generalized model, combined with variable binding strengths, give rise to a space in which evolution can optimize binding strengths in order to maximize the probability that critical assembly steps occur in the right order for a desirable phenotype.

### Baseline strength prediction

As mutations accumulate over the course of evolution, interaction binding strengths are unlikely to remain static. Predicting how binding strengths will evolve over time in a simplistic limit provides a comparative reference when examining evolution simulations. Several assumptions help reduce the mathematical complexity of the prediction, including

- no direct fitness advantage for stronger interactions
- falling below the critical strength is fatal
- infinite population
- only single mutations

Since selection can only operate on phenotypes, it is “blind” to the underlying genotypic details. Hence bonds present in the phenotype can be considered equal, justifying the lack of direct fitness advantage for interaction strengths. The remaining assumptions are fairly weak and satisfied by any reasonable choice of simulation parameters. These assumptions and the mutation-selection dynamics can be framed as a Markov process, giving both transient and steady-state expectations for the evolving interaction strengths. Details on this Markov process and calculating its expectations are in S2 Appendix.

### Simulated evolution

Interactions can be categorized on two distinct levels: phenotype and interaction topology. Selection acts on phenotypes, and so evolutionary dynamics may differ between phenotypes. Interaction topology can be characterized using two properties: The first is whether an interaction is inter-or intra-subunit, while the second is if either binding site in the interaction are involved in other interactions or if they are unique. Classifying interactions in this way allows different dynamics to be isolated, revealing the underlying causes.

Fig 4 displays the evolution of interaction strengths in the partial system. The three deterministic phenotypes (top left, bottom left, bottom middle) have similar behaviour, all approximately following the transient expectations of the Markov process (dotted black line), regardless of ancestral phenotype or interaction topology. Conversely, the three nondeterministic phenotypes (top middle, top right, bottom right) diverge from the expectations of the Markov process, with long-term interaction strengths being driven both above and below the Markov values. Notably, one interaction in the nondeterministic 16-mer does follow the Markov prediction, because it does not matter whether this particular assembly step occurs first or last.

**Fig 4.**
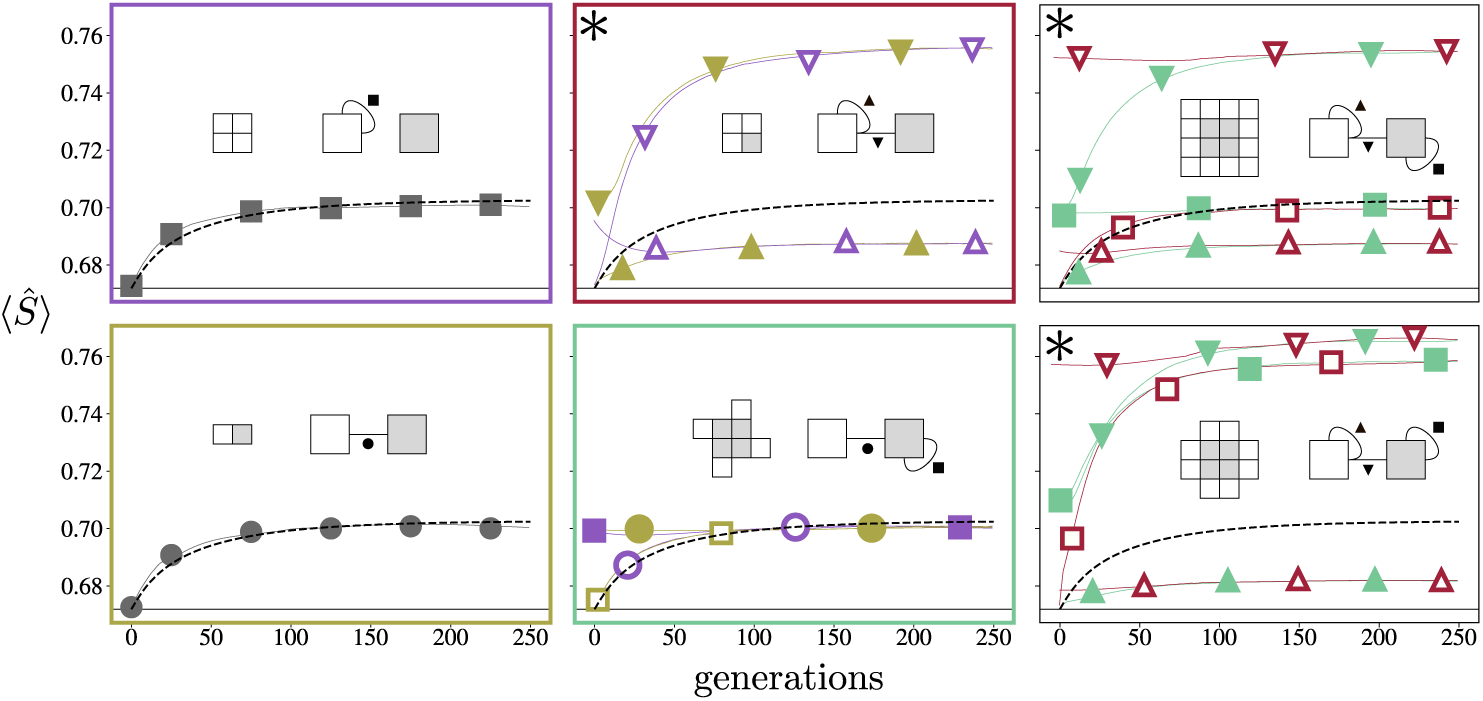
Binding strength evolutions. Each box corresponds to a different phenotype, with marker styles indicating interaction topology. Line colours (online) match the box colour of the direct ancestor, with “open” markers (print) indicating the ancestor is from an upper panel. Individual simulations are noisy, but averaging over many simulations yields stable trends. Black dashed lines in the panels are from the Markov prediction. The ∗ again indicates the three nondeterministic assemblies. Interface strengths in deterministic assemblies evolve predictably, while nondeterministic assemblies diverge rapidly.

### Selective ordering

In all three nondeterministic phenotypes the nondeterminism originates from the multiple possible orderings of individual assembly steps. Greater determinism can therefore be achieved by making sure that certain assembly steps occur earlier than others, by increasing the strength of the corresponding interactions. This is precisely what the evolutionary algorithm achieved through selection, with interactions strengthening or weakening across evolution to optimize determinism.

Such an effect is only observed in cases with steric nondeterminism, where for example the heterotetramer can be assembled with near 100% determinism if inter-subunit interaction binds much sooner than the intra-subunit interaction. Similar selective pressure for determinism drives the interaction strengths for the other nondeterministic assemblies.

### Universality

The choice of parameters, like nondeterminism punishment *γ* and “temperature” *T*, only have qualitative significance. Provided there is some fitness benefit to being more deterministic (*γ* > 1) and stronger interactions bind preferentially (*T* > 0), then the same patterns of behaviour are observed across a range of parameters. Exact values of the steady states vary intuitively with the choice of parameters, but the behaviour is near universal (see S1 Fig for more details).

## Evolutionary pathways

In the steady-state limit of the evolutionary simulations, mutation and selection effectively eliminate any trace of ancestry in the interface strengths. The steady state properties of interaction strengths depend only on the current phenotype. However, shortly after a new shape has evolved, it is possible to deduce ancestry from interface strengths. In the case of the 12-mer and the 16-mer, where we have one nondeterministic ancestor and one deterministic one, this is obvious as the interface strength distributions of the two ancestors differ considerably. As a result the two alternative ancestries for each of these two polyominoes can be clearly distinguished by bond strengths up to about 50 generations.

But even where we have deterministic ancestors, namely for the octomer and the heterotetramer, we notice that at the earliest time points the interface that is also present in the ancestor is stronger than the interface that is absent in the ancestor. This latter observation mirrors results found in real protein complexes, where the ordering of interface strengths often reflects the order of evolution, with the strongest interface as the oldest [10].

### Phenotype phase space

Deterministic assemblies, by definition, always produce the same polyominoes. On the other hand, nondeterministic assemblies can produce polyominoes with different frequencies due to the inherent stochasticity of the assembly process. In the limit of infinite repeated assemblies, these polyomino frequencies become deterministic and can be calculated *a priori*. The frequencies can be represented in a “phase space” for a set of nondeterministic interaction topologies. The ratio of interaction strengths provide the coordinates for the phase space.

These phase spaces can be calculated through a decision tree of assembly steps. Each branch in the decision tree is new binding step during assembly, and is weighted by the strength of that step’s interaction normalized by all possible step strengths. So the the final result does not depend on absolute values of interaction strengths, but rather ratios of the competing interaction strengths. The dimensionality of the phase space depends on how many competing interactions there are.

These trees quickly reach unusable levels of complexity due to exponential branching. Heuristics can eliminate many terms in the final expressions, identifying steps which are indistinguishable or deterministic. The decision tree calculation for a heterotetramer can be found in Fig 5.

**Fig 5.**
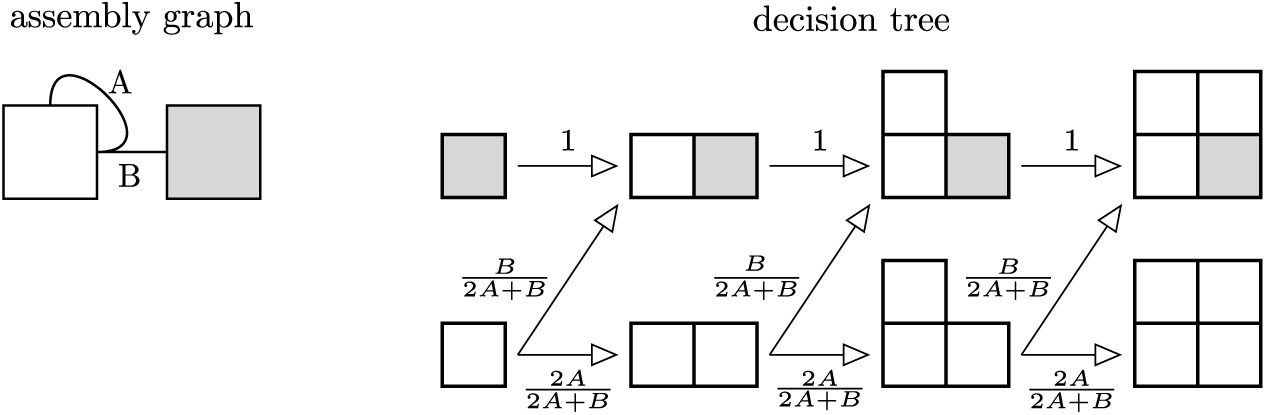
Decision tree for heterotetramer. The assembly graph has interaction strengths *A* and *B*. Each seed is a starting point for the decision tree, incrementally progressing until assembly terminates. In this situation, once a gray subunit is placed, assembly deterministically ends with the heterotetramer, rendering further branching unnecessary. The lower branchings have an extra weighting factor of two, due to two indistinguishable assembly steps.

### Simulated pathways

Phenotype transitions in a population are difficult to define precisely, so two general forms, fixations and failures, are introduced. Fixating transitions are those contained in any evolution history spanning the duration of the simulation, indicating they were beneficial transitions. Failures on the other hand, are transitions that quickly go extinct despite having higher fitness potential. Not all transitions fall within these two groups, but the remainder are artifacts of finite population size and can be explicitly ignored.

The success rate of transitions does not only depend on the properties of the descendant, but also depend on immediate ancestry, as shown in Fig 6 (a). Transitions to the heterotetramer for example, have very different success rates coming from the dimer or homotetramer, despite their qualitative similarity. The resolution to this apparent discrepancy is understanding the connection between a transition’s success rate and its location in the descendant’s phase space. Critically, the average location in this phase space can be predicted based on the ancestor’s steady state behaviour. The location in phase space in turn provides the level of nondeterminism and thus estimations on success rate, seen in Fig 6 (b) and (c).

**Fig 6.**
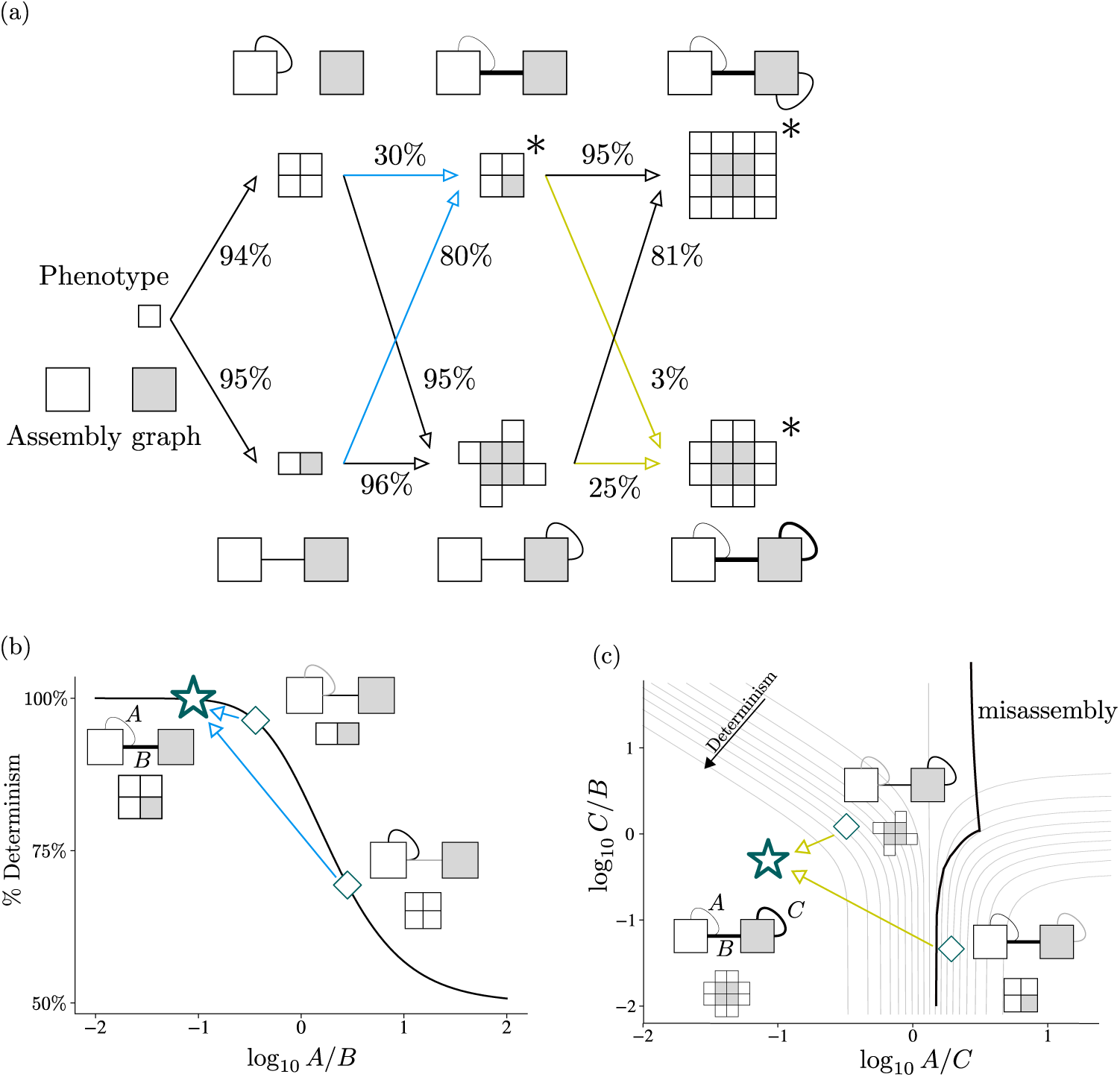
Phenotype transition success and ancestry. (a) Transitions to deterministic assemblies have high success, tending to perfect in an infinite population. Conversely, transitions to nondeterministic assemblies (marked with *) typically have less success. Transition rates between nondeterministic assemblies vary considerably, due to the varying overlap between the interfaces of an ancestor and the stronger interfaces of the descendant. Interaction strength is indicated by line thickness. The transition locations in phase space of ancestors are shown for the heterotetramer and 12-mer in (b) and (c) respectively. (b) For transitions from both the dimer and homotetramer, one bond has been strengthened through evolution (black) and one is new and at minimum value (gray). Compared to the evolutionary equilibrium of the heterotetramer, the dimer has a much more favorable ratio of strengths than the homotetramer, as indicated by its closer position in phase space. Likewise in (c), the evolutionary equilibrium of the 8-mer has much more similar ratios of interaction strength to the 12-mer than the heterotetramer has. In addition to the heterotetramer being further down the determinism gradient, it more frequently misassembles the phenotype, lowering its transition success even further.

There are 3 pairs of transitions that are interesting to examine: those to the heterotetramer, 16-mer, and 12-mer. For the heterotetramer, as can be seen from its phase space in Fig 6 (b), the assembly is most deterministic if the inter-subunit interaction is significantly stronger than the intra-subunit interaction. The average transition from the dimer is much closer to this constraint than the average transition from the homotetramer, and this is reflected in the success rates (80% compared to 30% respectively).

As noted earlier, one interaction in the 16-mer does not compete in assembly order, and the 16-mer actually shares the same decision tree as the heterotetramer. Trivially the heterotetramer will evolve to its own optimal interface strength ratio, and thus transition in the optimal location for the 16-mer. This is reflected with its effectively deterministic success rate (95%). The 8-mer is effectively the dimer once discounting the non-competing interaction, and transitions in the same region with similar successes of about 80%.

The 12-mer phase space is more complicated, with three competing interactions and three possible polyominoes, although only the 12-mer and “misassmbled” states are of interest here. Analogous to before, the 8-mer transitions higher on the determinism gradient and thus is more successful than the heterotetramer. However, these assembly graphs can misassemble more often than they assemble the 12-mer, and thus produce an unfit phenotype. The average transition for the heterotetramer is fatal, because it occurs in the misassembly region, seen in Fig 6 (c). Stochastic fluctuations can shift the individual transition locations, but such an event is a “second-order probability”. As such, the heterotetramer to 12-mer is strongly constrained, and has a meager 3% success rate.

More exact calculations can be done to predict transition success rates from phase space locations, but these depend explicitly on the nondeterminism parameter *γ* and how much more fit each descendant is. However, as before, the behaviour is qualitatively near-universal. These transitions are taken directly from the simulations displayed before, again with parameters chosen to highlight these dynamics clearly.

## Discussion

### Ordered assembly

The time ordering of assembly steps in proteins is integral to the correct assembly of the protein structure. This holds true on many length scales of assembly, with cotranslational protein folding able to induce misassembly [15] all the way up to final quaternary structure as examined here. Experimental methods for devising binding strengths are still being developed [16], with an *in silico* approach recently introduced focusing on multimeric complexes [17].

One notable result was that given an equal rate of mutation, deterministic and nondeterministic assemblies adapted at different rates. The peak observed rate of binding strength increase in the 12-mer was approximately triple the rate in deterministic assemblies. Such an observation is fairly intuitive, as mutations which alter binding strength correctly or incorrectly are more strongly selected or purified respectively in the nondeterministic assemblies. This is in good agreement with the observation that unstable proteins adapt more quickly [18].

Binding strengths that deviate from neutral expectations do so to optimize determinism, assembling a core of the final structure as quickly as possible before adding further, peripheral elements. This evolutionary selection for a particular assembly pathway has an equivalent in real protein complexes, in which gene fusions are a way of cementing particular assembly order under evolutionary selection pressure in order to minimize the risk of misassembly [11].

### Model implications

Generalizing the binding sites from integers to binary strings provides a range of benefits. The number of binding site configurations is now fixed by a physically meaningful parameter and is exponentially large. Previous models frequently had identical binding sites at multiple locations, which is very unlikely in real proteins, whereas now repeated binding sites are vanishingly rare. Additionally, interaction rules in the integer model have trivial transitivity relations: Maintaining the notation of ↔ for interactions, that is to say for sites *A, B, C* that

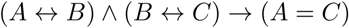

However, the generalized model does not require the above relation to be true, with knowledge of one interaction having little bearing on other interactions sharing a binding site. That it is to say for sites *D, E, F, G* that

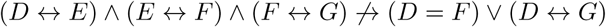

This allows more complex interaction patterns to form, but also allows different binding sites to produce the same interaction behaviour, as seen in Fig 7. In addition, sites can self-interact, interact with another binding site, or both, like sites *D* and *E* supporting the interactions *D* ↔ *E* and *E* ↔ *E*.

**Fig 7.**
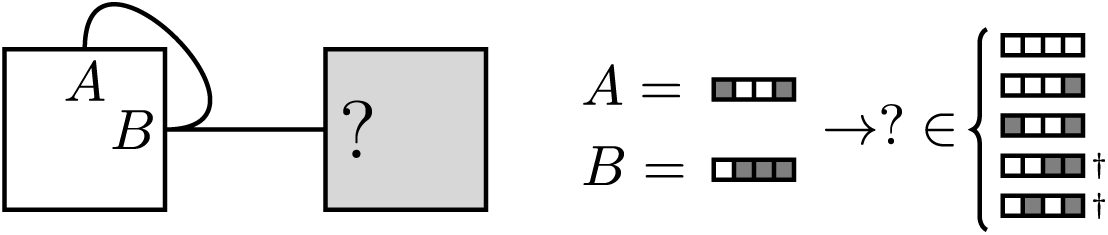
Generalized interactions are not always transitive. In the generalized model, knowledge of one interaction does not fix the binding sites of another related interaction. Earlier in the nondeterministic case in Fig 1, this assembly graph had *A* = 1, *B* = 2 fixing ? = 1. Here, choosing binding sites *A* and *B* still leaves 5 possibilities for ?. The possibilities marked with † self-interact, and so would technically add an interaction to the assembly graph.

Usefully, the generalized interactions are a superset of the integer model, and so any previous results could be trivially recovered by choosing *Ŝ*_*c*_ = 1 (up to relabeling binding sites). While the generalized model is still a very abstract representation of biological self-assembly, the binary interfaces add physical realism and layered complexity to an already promising model.

## Extensions

Phenotype plasticity is another feature that is naturally introduced by the generalized model. By incorporating a dynamic fitness landscape, one that alternatively favors two (or more) phenotypes, the interaction strengths can continuously adapt to remain optimal, shown in Fig 8. The ability to modify a phenotype in a controllable manner, minimizing nondeterminism, is a huge advantage to survival. If a conformational change of a protein, in response to an environmental change or other external conditions, altered its binding strengths, it could quickly shift phenotypes.

**Fig 8.**
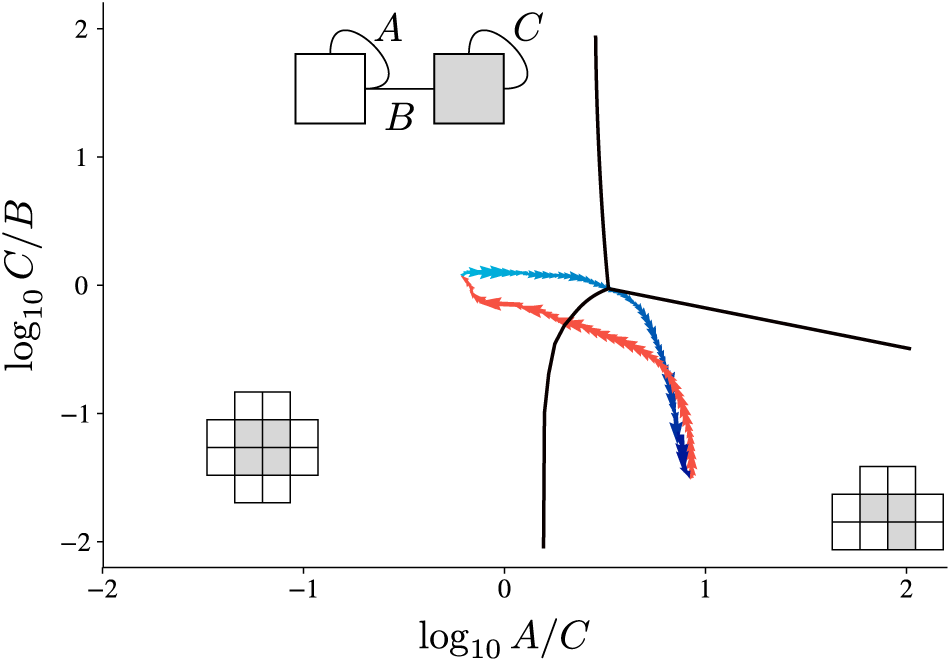
Interaction strengths can adapt to changing fitness landscapes. Periodically alternating the fitness landscape produces cyclic behaviour in interface strengths. Despite starting from a range of initial conditions, all simulations eventually converge to the optimal path to transition between the 10-mer and 12-mer and back. The change in fitness landscape is indicated by the red or blue colours, with arrows indicating the direction of flow. Both phenotypes are produced with the same three interactions; it is only the relative ordering of interaction strength that matters. A breakdown of each fitness landscape and local gradients can be seen in S2 Fig.

Since changing interaction strengths can occur much quicker than creating new interactions, this plasticity allows adaptions that would otherwise be potentially too slow to survive. The relationship between conformational changes and their impact on evolution is uncertain, but it has been suggested that this behaviour can impose strong constraints on sequence evolution [19, 20]. Moreover, adding and removing interactions, rather than just reprioritizing them, exposes the assemblies to intermediate states and greater risk of negative outcomes [21].

## Conclusion

Polyomino self-assembly models using integers as binding sites have demonstrated the value of abstract self-assembly models for the study of self-assembly phenomena and genotype-phenotype maps [2–4].

Generalizing the binding interfaces using binary subsites as outlined in this paper retains tractability while expanding applicability to more complex biological research questions. In particular, modeling the evolution of interaction strengths provides qualitative insights beyond the reach of previous polyomino studies.

With a few justifiable assumptions, analytic predictions of the interaction strengths in the absence of selection pressures can be found, which show strong agreement with simulations. Significant divergences from this prediction are observed in nondeterministic assemblies where time-ordering is important, and the interaction strengths are therefore under selection. This selection pressure drives these interactions to strengthen or weaken, and thus bind earlier or later in the assemble, to optimize the determinism. Certain interaction strength orderings are more suitable for transitioning to descendant phenotypes, and so can be used to statistically reconstruct evolutionary pathways.

Several observations from experimental studies have been recovered by this model, as well as suggesting that nondeterminism in the Polyomino model provides an interesting framework for the study of protein misassembly. Many further avenues are imaginable that build on such investigations of nondeterminism, including gene duplication, phenotype plasticity, and more complex genotype-phenotype mappings.

## Methods

A full implementation of the self-assembly algorithms, evolutionary dynamics, and phylogenetic analysis written by the authors can be found online [22].

## Evolution

As outlined earlier, evolution was modeled with asexual reproduction of haploids encoding two subunits (total of 8 binding sites per genotype). Binding site lengths were *L*_*I*_ = 64 and the critical strength was taken as *Ŝ*_*c*_ = .671875. Genotypes were initialized randomly, with the constraint that there were no interactions. Assembly could begin with either subunit as the seed, although monomers were ignored due to their trivial contribution.

A population of 250 individuals evolved for 1000 generations, with each genotype being assembled 25 times. Each binary subsite had a fixed probability to flip, such that the entire genotype had mutations that were binomially distributed with mean *µ* = 1. The temperature was set to *T* = 25, while the nondeterminism punishment was *γ* = 5.

An individuals fitness was calculated as 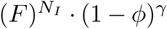, where *F* is the fitness jump between higher order assembly graphs, *N*_*I*_ is the number of interactions in an assembly graph, and *ϕ* is the nondeterminism fraction for that particular set of assemblies. The fitness jump was set to 5 to balance the strong nondeterminism punishment.

Similar results were achieved with different binding site lengths, critical strengths, fitness functions, etc. Likewise, mutation rate, population size, and other simulation dynamics all displayed the same qualitative behaviour. The parameters used in these results offered good fidelity and reasonable computation timescales, but were otherwise arbitrary.

### Phylogenetic tracking

With asexual reproduction, new interactions or new phenotypes can be traced directly to unique mutation events. The descendants of these individuals can be tracked for separate evolutionary histories. By recording the assembly graphs, phenotypes, and reproducing individuals at every generation, the ancestral information can be entirely reconstructed.

### Dynamic landscapes

The bulk of the results were attained with static fitnesses, but Fig 8 had two distinct fitness landscapes alternating periodically. Here, an individuals fitness was taken as the *l*_1_ norm of fitnesses in both the 10-mer and 12-mer landscapes at that generation. The rate at which the landscapes varied smoothly was only of qualitative importance, provided that timescale was significantly greater than the mutation timescale.

## Supporting information

**Fig S1.**
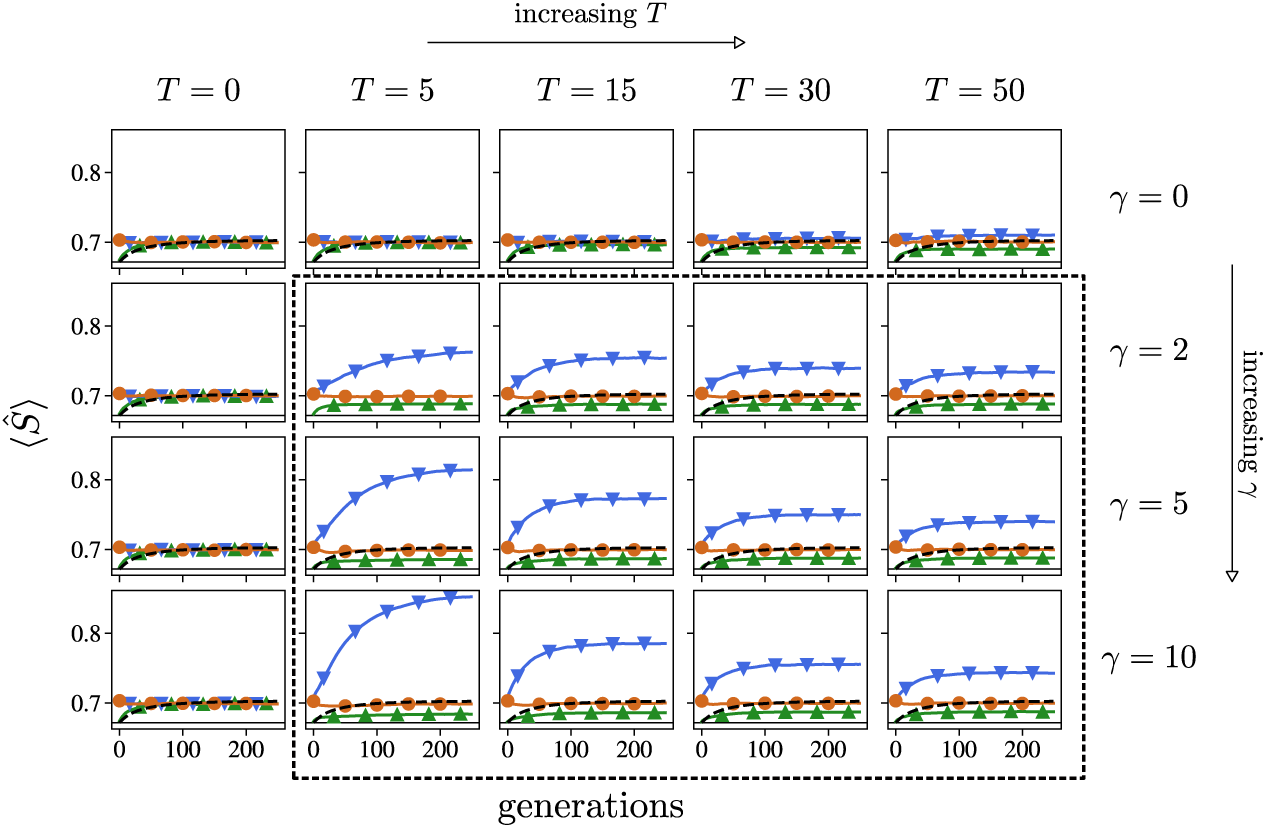
Binding strength evolutions are qualitatively universal. For all values of *T* > 0 and *γ* > 1 (in dashed box), where the parameter space enabling stronger bonds to optimize determinism, the same qualitative observations hold. The equilibrium values of interaction strength do depend on the selective pressure and temperature, but vary intuitively.

**Fig S2.**
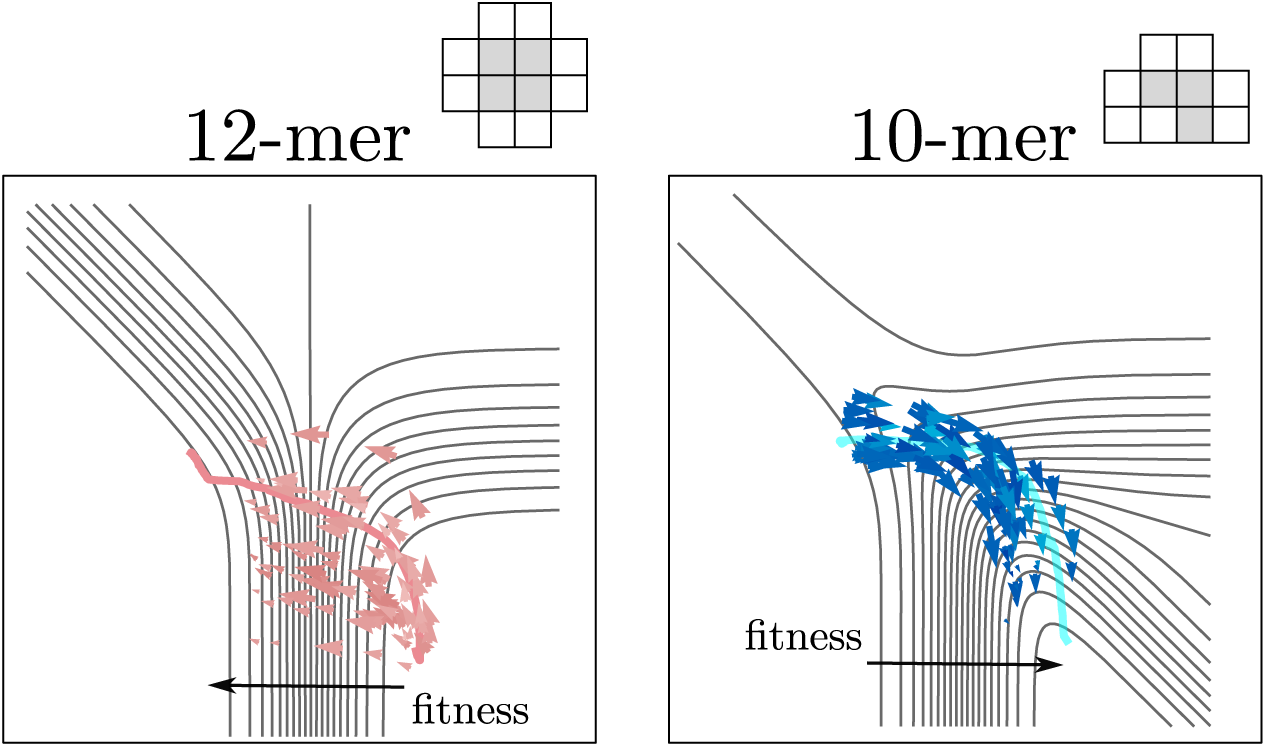
Interaction strength adaptation follows determinism gradients. After switching the rewarded phenotype in the fitness landscape, average trajectories closely follow the determinism gradient of the relevant phenotype. Some trajectories switching from the 10-mer to the 12-mer (red) follow local gradients, increasing the *C/B* ratio first, as opposed to the more global optimum of lowering the *A/C* ratio. However, both paths tend to the same steady-state region of phase space.

**S1 Appendix. Polyomino comparison.**

**S2 Appendix. Markov evolution.**

